# TXO: Transcription-Only genetic circuits as a novel cell-free approach for Synthetic Biology

**DOI:** 10.1101/826230

**Authors:** Felipe A. Millacura, Mengxi Li, Marcos Valenzuela-Ortega, Christopher E. French

## Abstract

While synthetic biology represents a promising approach to solve real-world problems, the use of genetically modified organisms is a cause of legal and environmental concerns. Cell-free systems have emerged as a possible solution but much work is needed to optimize their functionality and simplify their usage for Synthetic Biology. Here we present TXO, transcription-only genetic circuits, independent of translation or post-translation maturation. RNA aptamers are used as reaction output allowing the generation of fast, reliable and simple-to-design transcriptional units. TXO cell-free reactions and their possible applications are a promising new tool for fast and simple bench-to-market genetic circuit and biosensor applications.

## Introduction

Cells, including microbial, plant and mammalian cells, have been long engineered for responding to a plethora of environmental factors, benefiting from their intrinsic ability to process the entire cycle from the recognition of a specific target, to analysis of the information, and generation of an output signal [1]. However, cell-based systems present also unavoidable disadvantages during their usage, including the risk of releasing genetically modified organisms (GMOs) into the environment, accumulation of mutations and genetic instability, uncontrollable side-reactions within the cell, and issues with viability maintenance during long-term storage [2–5].

In order to solve these issues, cell-free systems (CFS), also known as transcription-translation (TX-TL) systems, have emerged as a promising alternative. CFS not only inherit most of the benefits from cell-based systems, but also avoid major challenges that they face. Typically CFS are based on non-living cell-extracts or reconstituted systems that by not presenting a living prospect solve problems related to biosafety, cross-reactivity and long-term functionality maintenance. CFS are rather robust to factors that pose cellular stress, including sensitivity to toxic pollutants and to the typical “overload problem” observed when foreign “genetic circuits” are introduced into the cell [6].

TX-TL systems share with cells a problem hindering their usage in real-time detection. Response time in TX-TL systems may need a long time for transcription, translation and post-translational maturation to occur, in order to finally produce a detectable signal. For instance, Pardee and colleagues [7] developed a cell-free paper toehold-switch sensor for detection of the Zika virus that takes hours for the final visual results to come out [6]. Paper-based cell-free sensors that detected heavy metals and gamma-hydroxybutyrate by expressing sfGFP needed at least one hour for producing a measurable response [8]. Considering the need for in-situ field analysis, an improvement of cell-free systems response time is necessary [9, 10].

Since CFS are open and easily modifiable systems [11], time-consuming translation and post-translational modification can be removed, allowing novel approaches with faster signal generation. Alternative output signals independent of translation and post-translational modification can provide the system with a simpler composition, lower resource requirements and, most importantly, a shorter response time. Here, we introduce a transcription-only (TXO) cell-free system using RNA aptamers as output signals rather than translated proteins.

Fluorescent RNA aptamers are short ribonucleotide sequences that become fluorescent when bound to specific fluorophores. RNA aptamers, such as Spinach [12], iSpinach [13] and Broccoli [14] have shown excellent properties for generating bright and stable fluorescence. In the past few years, RNA aptamers have been widely applied in live-cell imaging [15] and biosensing [16]. However, these previously reported aptamer-based sensors acted as both sensing elements (binding to target molecules) and signal generator at the same time [17,18], requiring aptamers specifically designed for each target. In TXO systems the functions for input detection (sensing), processing (analysis) and output generation (reporting) are decoupled, creating modular systems where aptamers can be used as universal signal generators. To exemplify the versatility of such system, here we used a simple cell-free extract from the microorganism *Cupriavidus metallidurans* CH34, which carries most of the necessary heavy-metal (e.g., Hg^+^, Pb^2+^, Cu^2+^, Zn^2+^, Ni^2+^, among others) responsive transcription factors and the RNA polymerase which they control [19]. Since all necessary transcription factors are provided directly by the cell-free extract only the synthetic transcription unit must be changed to detect these different metal ions. Combining of the easy to design TXO system, with the high sensitivity of the sensing elements and the fast output signal generation provided by RNA aptamers, TXO is a promising tool for accelerating bench-to-market Synthetic Biology, most specifically where in-situ, simple to design and rapid applications are needed.

## Materials and Methods

### Reagents and culture conditions

Fluorophores DFHBI (Tocris #5609), DFHO (Tocris #6434), DMABI and 2-HBI (provided by Dr. Jaffrey’s laboratory at Cornell University) were dissolved in DMSO (Fisher Scientific #BP231) to prepare 2 mM stock solutions (for further information see Table S1). Our optimised transcription/detection buffer (OTDB) was prepared as a 10X stock as follows: Tris base 400 mM pH 7.5 adjusted with HCl (Sigma-Aldrich, #T1503); MgCl_2_ 6(H_2_O) 60 mM (Sigma-Aldrich, #M2670); DTT 100 mM (Melford, #MB1015) and Spermidine 20 mM (Alfa Aesar, #A19096.03). Metal-ion solutions were prepared from soluble salts of analytical grade: CuSO_4_·5H_2_O (Sigma-Aldrich #C8027), Arsenite NaAsO_2_ (Sigma-Aldrich #202673), Pb(NO_3_)_2_ (Sigma-Aldrich #228621), HgCl_2_ (Sigma-Aldrich #215465), ZnCl_2_ (Sigma Aldrich #229997) and NiCl_2_ 6H_2_O (Fisher Scientific #AC27051) in double-deionised water and filter-sterilised before use.

### Strains and cell-free extract preparation

*C. metallidurans* CH34 cells were cultivated on a Tris-buffered mineral medium (MM284) [20] supplemented with succinate 0.4 %w/v, at 30*°*C with constant shaking at 180 RPM. *Escherichia coli* JM109 cells were grown on Luria Broth (LB) at 37*°*C. For cell-extract preparation, cells were collected at middle exponential phase (OD 0.6-0.8) by centrifugation at 3,900 × *g* and 4*°*C for 10 min. Cells were resuspended in one tenth of the centrifuged volume in a non-ionic lysis buffer, which contains Triton X-100 0.1%v/v (Sigma-Aldrich #T9284), Tris base (40 mM) adjusted to pH 7.5 with HCl and lysozyme 50 mg/mL (Sigma-Aldrich #L6876) for 30 min at 4*°*C. The lysed extract was centrifuged at 12,000 *g*, 4*°*C for 20 min and the supernatant was immediately used or saved at -80*°*C for later use.

### Construction of RNA aptamers

Aptamer sequences (see Table 1) protected within the F30 scaffold (upper and lower sections) were synthesised as 120 bp oligonucleotide sequences (Sigma-Aldrich), and then extended using the 5 bp shared on the 3’ of each oligonucleotide for the annealing (see Figure 1 and Table S2) A PCR reaction adding a T7 promoter/terminator pair on each aptamer (see Table 1) was carried out using Q5 High-Fidelity DNA Polymerase (New England Biolabs Inc.(NEB) #M0491) and the following protocol: 95*°*C for 5 min (1 cycle), 95*°*C for 20 s, 54*°*C for 10s and 72*°*C for 20 s (35 cycles), with a final elongation at 72*°*C for 5 min. Amplification was analysed in a 1.5 % (w/v) agarose gel and purified from the PCR reaction using the QIAquick PCR Purification Kit (Qiagen, #28104), and subsequently cloned in to the pMini-T plasmid using the PCR Cloning Kit (NEB #E1202). Plasmids were extracted from *E.coli* JM109 transformed cells using the QIAprep Spin Miniprep Kit (Qiagen) and sequence verified by Sanger sequencing using the BigDye Terminator v3.1 Cycle Sequencing Kit (Thermo Fisher Scientific) and an Applied Biosystems 3730XL DNA Analyzer (Edinburgh Genomics).

**Table 1.** RNA Aptamers and their respective characteristics

| Aptamer | Emitted Colour | Size (bp) | Ex/Em (nm) | Fluorophore | Reference |
| --- | --- | --- | --- | --- | --- |
| Broccoli | Green | 47 | 470/530 | DFHBI | [14] |
| Broccoli2X | Green | 157 | 470/530 | DFHBI | [21] |
| Spinach2 | Green | 95 | 480/520 | DFHBI | [12] |
| iSpinach* | Green | 78 | 480/520 | DFHBI | [13] |
| Corn | Orange | 45 | 480/520 | DFHO | [22] |
| 2-4** | Cyan | 100 | 400/460 | DMABI | [12] |
| 17-3*** | Yellow | 101 | 400/550 | DMABI | [12] |
| 6-8**** | Red | 101 | 400/600 | 2-HBI | [12] |

**Figure 1.**
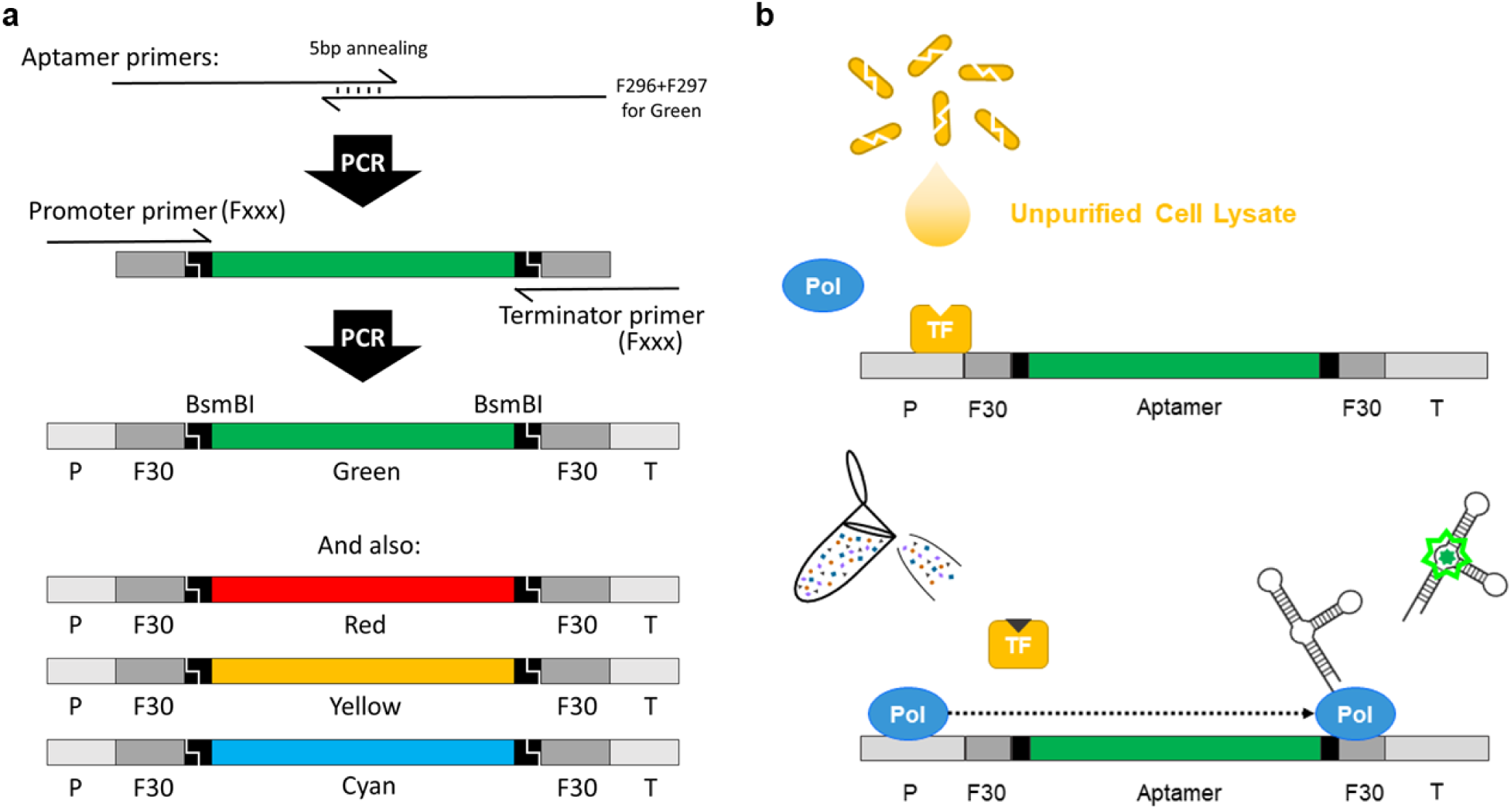
Schematic diagrams of TXO cell free system with minimal requirements. **(a)** Sequence and production process of diverse RNA aptamers via PCR, showing addition of BsmBI sites for further Golden Gate manipulation. **(b)** In a TXO system there are minimal requirements such as an inducer, moderator/repressor, RNApol and a final RNA output herewith represented as a fluorescent aptamer.

### Biosensor transcriptional unit design

Diverse inducible promoters and a T7 terminator were added at the ends of the aptamer F30::iSpinach via PCR [13]. The PCR was carried out using Phusion High-Fidelity DNA Polymerase (NEB #M0530) and oligonucleotides containing the promoters from clusters *copAB, arsBCD* from *E. coli* MG1655 and *merTPAD, pbrAB, czcN, cnrH* from *C. metallidurans* CH34 (see Table S2). Amplified products were analysed and cloned for replication as described above.

### RNA aptamer expression and detection

Some previous protocols for *in vitro* transcription of fluorescent RNA aptamers [12, 13] include purification and concentration steps that complicate the use of aptamers for real-time detection. A combination of both main steps, expression and detection, was achieved at room temperature (25*°*C) by using the OTDB buffer. *In vitro* transcription and detection assays were carried out using OTDB 1X, NTPs (2 mM each), Fluorophores 10 *µ*M (see Table 1), NEB T7 RNA Polymerase (1.5 U/*µ*L), linear target DNA (650 nM), using 15 *µ*L total reaction volume in 384 Well Small Volume*™* LoBase Black Microplates (Greiner Bio-One). Fluorescence was measured using a FluoStar Omega (BMG Labtech) plate reader, 10 mm bandpass, using 20 flashes per well and an orbital averaging of 4 mm, with 5 seconds of orbital shaking prior to each measurement, and filters as close as possible to the wavelength specified in literature (see Table 1). Fluorescence was thereafter imaged using a Safe Imager*™* Blue-Light Transilluminator (Invitrogen) with an amber filter unit and a digital camera (Sony, DSC-RX100).

### Heavy metal detection by TXO assays

Transcriptional regulators MerR (Q5NUU7), PbrR (Q58AJ5), CzcS (Q44007)/CzcR (Q44006), CnrH (P37978), ArsR (Q1LCN6) and CueR (A0A2L0XBJ2) were provided into the working reaction directly in the form of cell extract from *C. metallidurans* CH34 cells (30 %v/v). Reactions were supplemented with increasing concentrations (0.3 *µ*M, 2 *µ*M and 7 *µ*M) of Hg^2+^, Cu^2+^, Co^2+^, Ni^2+^, Pb^2+^ and As^3+^. Cell-free metal contact induction was performed in 384 Well Small Volume*™* LoBase Black Microplates (Greiner Bio-One) at 25*°*C, using DFHBI (10 *µ*M) as fluorophore, and analysed in a FluoStar Omega (BMG Labtech) plate reader as described above.

## Results

### Fluorogenic RNA aptamers and TXO systems

Commonly, cell-free systems are associated with the use of TX-TL systems instead of living cells for the production of valuable compounds or the generation of translational biosensors. Their dependence on translation, however, increases the complexity and cost of the minimal reaction mixture needed for them to work. In order to generate an alternative approach, we evaluated the use of multiple fluorescent RNA aptamers as possible output signals for the generation of transcription-only cell-free systems.

Eight previously characterised fluorescent aptamers were initially screened (see Table 1. These were synthesised as 120 bp oligonucleotides and amplified in an overlapping PCR using the 5 bp that they share at their 3’ end (see Material and Methods, and Figure 1 and Table S2). Additionally, unprotected aptamers (Green, Cyan, Yellow, Red, Corn) were inserted within the F30 RNA scaffold, an engineered version of the naturally occurring phi-29 viral RNA three-way junction motif, not cleaved by cellular nucleases [21]. Additionally, BsmBI sites were added at both ends of each aptamer to allow insertion or replacement of the RNA aptamers via Golden Gate assembly if required in future applications (see Figure 1). RNA expression and fluorescence detection were performed at room temperature (25*°*C) in order to simulate field conditions as closely as possible. Furthermore, in order to simplify their use we combined the expression and detection steps by using the novel OTDB buffer (see Materials and Methods), avoiding purification steps needed in some previous cell-free publications. Aptamers were selected primarily by difference in the colour emitted (green, red, orange, yellow, cyan), stability once protected within the F30 scaffold [21], length, fluorophore needed, and expression level under our previously mentioned experimental conditions (see Table 1).

The linear PCR-generated fragments can be immediately used for *in vitro* transcription and fluorescence detection. We tested the expression of nine different aptamers with five different fluorophores using a plate reader and an inexpensive blue light/red filter system (see Figure 2). While previous studies regarded aptamers to be specific for a fluorophore (see Table 1), we have found that cross-reactivity is much higher. As shown in Figure 2a, Spinach, Broccoli and their derivatives, engineered to bind DFHBI, were found to fluoresce with DMABI and DFHO with their respective colours. Broccoli2x was the most unspecific even increasing the fluorescence of ROX, which we were using as positive fluorescence control. On the other hand, DFHO was the most promiscuous fluorophore, which also displayed background fluorescence with the negative control. The aptamer fluorescence appeared very quickly after induction of transcription; after just a few minutes of incubation at room temperature, there was a significant difference between some aptamers and the negative control (see Figure 2b). However, the fluorescent signal kept increasing for 100 minutes (Figure 2b), indicating that the RNA polymerase remained active and the NTP pool was not exhausted for this period. Additionally, all F30::aptamers presented high stability over time when left on the bench at room temperature and without further protection. Aptamer-fluorophore complexes were stable for weeks without bleaching or loss of fluorescence (See Figure S1).

**Figure 2.**
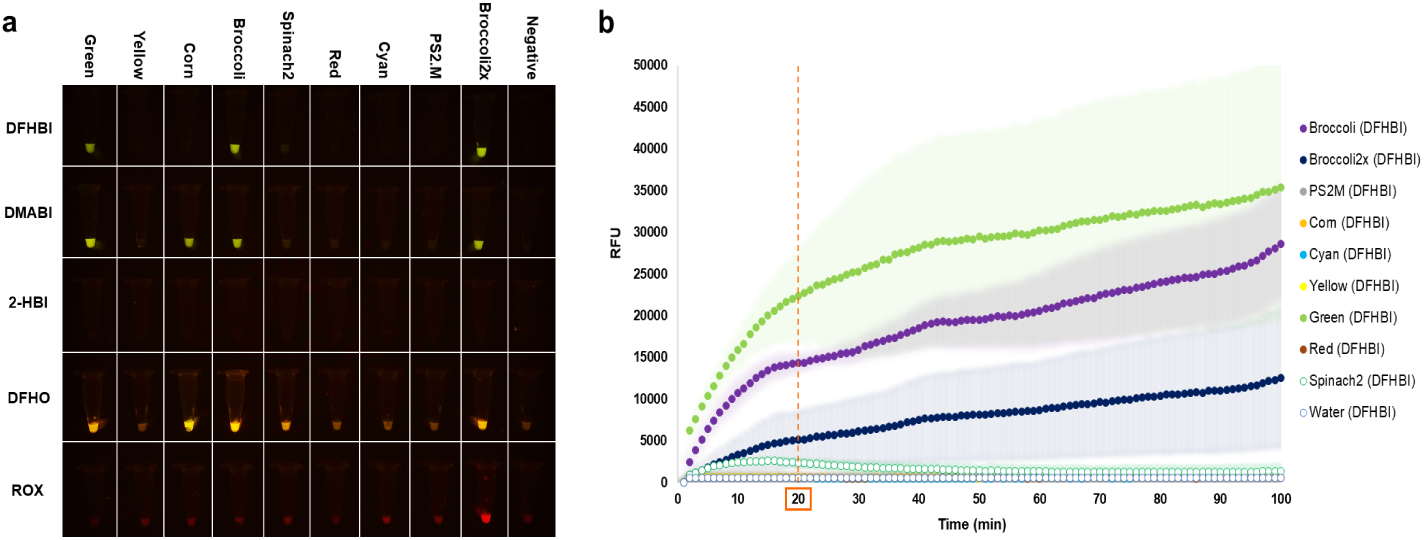
Fluorescent RNA aptamers expression. **(a)** Aptamers showing different colours under a blue light/red filter system after plate reader fluorescent measurement (16.5 h) and being observed by naked eye without the need for further equipment. A longer exposed image can be found in Supplementary Figure S1. **(b)** Expression of diverse RNA aptamers over time detected at 480/520 nm. Coloured shades represent the variability (MAD) of three independent tests.

### TXO cell-free biosensors

Short aptamers such as the Green aptamer allow incorporation into transcriptional units with ease and simplicity (see Materials and Methods). Considering the stability and optimization of the Green aptamer (F30::iSpinach) for expression under *in vitro* conditions, further experiments were performed with this aptamer. The Green aptamer (F30::iSpinach) was amplified via PCR under the control of diverse heavy metal inducible promoters and the T7 polymersase promoter as control (P*t7*, P*cue*, P*ars*, P*mer*, P*pbr*, P*czc*, P*cnr*, see Material and Methods). Linear PCR products were used in reactions with different heavy metal concentrations (5 *µ*M, 25 *µ*M and 100 *µ*M) with transcription factors and native RNA polymerase supplied from cell lysates (see Figure 3). Using our TXO cell free system concentrations as low as 2 *µ*M were detected for lead, mercury, zinc, and arsenite. Nickel, on the other hand, was detected at 0.3 *µ*M. All of them reached a 2 fold-change in fluorescence after just 20 min of initiation of the transcription reaction (see Figure 3) which kept increasing over time reaching after three hours values up to 10-times higher (see Figure S2).

**Figure 3.**
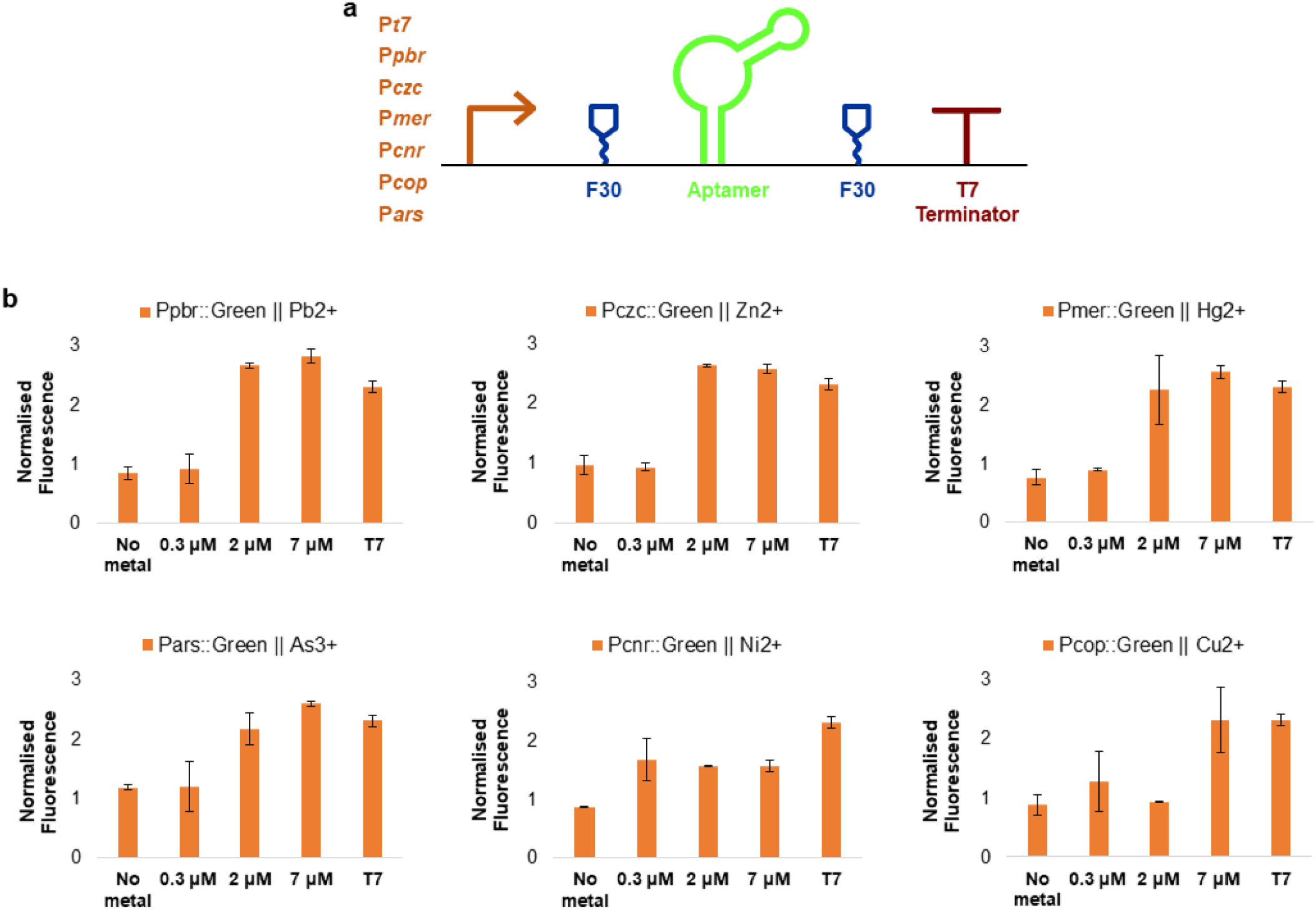
Heavy metal detection by TXO Cell-free biosensors. Detection of Pb^2+^, Hg^2+^, Zn^2+^, Ni^2+^, Cu^2+^ and As^3+^ using the TXO reaction Cell-free system and the Green RNA aptamer cloned with each specified promoter (P*pbr*, P*mer*, P*cnr*, P*czc*, P*cop*, and P*ars*). Constructs were induced using increasing concentrations of the specified heavy-metal. Excitation/Emission of 480/520 nm. Data points at 20 minutes; the time course can be found in Supplementary Figure S2. Error bars represent the variability (MAD) of three independent tests.

## Discussion

### Subsection heading

TXO systems maintain the advantages of cell-free TX-TL systems in terms of uniformity and avoidance of regulatory issues related to live GMOs, and further increase the simplicity and speed of response at the expense of reduced flexibility, since new proteins can not be generated within the system. Hence they are well suited to applications where simplicity and rapid response are paramount; for example, biosensors and diagnostics for use in field and point-of-care applications. Lack of translational capability means that reporters for system output must be in the form of nucleic acids. We tested eight previously described fluorescent aptamer-fluorophore systems and found that all could generate visible fluorescent responses in an optimized buffer system as a result of transcription with no further processing. Several of the aptamers showed the ability to bind a wider range of fluorophores than what has previously been reported; for example, Broccoli2X binds 4 out of the 5 fluorophores analysed (Figure 2a and S1), surprisingly including ROX (5-Carboxy-X-rhodamine N-succinimidyl ester) commonly used for basal line generation during routine fluorescent assays, such as qPCR. Using TXO systems fluorescent responses can be generated in a matter of minutes, as compared to an hour or more for reported TX-TL systems based on generation of fluorescent proteins or enzymes with chromogenic substrates [6, 23–27]. Thus we conclude that aptamer-fluorophore complexes are a suitable output for TXO synthetic biology systems.

A further advantage of TXO systems is simplicity, meaning that they can be very rapidly generated and tested, even more so than TX-TL systems [7]. The aptamer-based reporter genes are small enough that they can be easily generated by annealing and extension of oligonucleotides (see Figure 1). PCR was used to add metal-responsive promoters to DNA encoding the F30::iSpinach reporter (‘Green’), and the PCR products were used directly for assays, with the necessary transcription factors being supplied by an unpurified lysate from *C. metallidurans* CH34. Fluorescent response to metals was observed within minutes, with detection limits in these unoptimized systems below 0.3 *µ*M (18 ppb) for nickel, and below 2 *µ*M for copper (127 ppb), zinc (131 ppb), arsenic(III) (150 ppb), mercury (403 ppb) and lead (414 ppb). No response was observed for arsenic(V), presumably because the system lacked the reducing power necessary to reduce it to arsenic(III), the form actually detected by the ArsR repressor (Data not shown). Further optimization should reduce these limits of detection. Previous works with RNA aptamers for metal sensing, DasGupta et al. mentioned the use of a truncated form of the Spinach aptamer as a Pb^2+^ sensor, but this could have lead to false results at concentrations *<*2 ppm due to a lack of sensitivity and a weak output response [30].

Since TXO systems are considerably simpler than TX-TL systems, with no requirement for ribosomes, tRNA, or associated enzymes, we expect that it will be possible to freeze-dry such systems for storage and distribution as previously reported for TX-TL systems [8, 28]. Fluorescent signals may be further strengthened using alternative fluorophores such as DHFBI-1T [29]. The visibly different colours, as well as different excitation and emission wavelengths for different aptamer-fluorophore complexes, raise the possibility of multiplexed assays as well as more complex approaches such as FRET and quenching. While this publication focuses on the use of aptamers as signal output in a TXO ststem, future research could exploit the advantages of translation independent systems introducing other known functions of RNA molecules. Examples include RNA-based molecule detection (e.g. using riboswitch aptamers) and signal processing (e.g. using RNA-RNA hybridisation networks or crRNA/gRNA) [30–35]. Possible improvements to the already existing technology could also imply the use of other RNA aptamers showing different properties, for instance, by improving the structure and conditions of other aptamers published in the original work by Paige *et al*, which emit fluorescence in different wavelength, i.e. red, yellow, cyan (see Table 1 and Figure 2) or by using ribozymes, such as PS2.M, allowing additional possible transcriptional outputs and broadening the possibilities of TXO systems.

Additionally, TXO systems can be made with no need for GMOs, using in vitro DNA manipulation and non-ionic detergent lysis of wild type ACDPI or Biosafety level 1 cells, plus non toxic chemicals and commercially available enzymes/proteins, so procedures could easily be done in high schools or even at home, leading to a democratization of synthetic biology without the worrying risks of people making bioweapons or causing ecological disasters. Considering that algorithms are simply a sequence of instructions, typically to solve a class of problems or perform a computation, TXO systems offer a step forward in terms of simplicity, ease of implementation, and rapid response, and may be widely useful in synthetic biology for speeding up and simplifying genetic circuit algorithm design and response.

## Acknowledgments

The authors acknowledge the valuable assistance of Dr. Samie R Jaffrey on providing diverse RNA aptamers and fluorophores, Dr. Michael Ryckelynck for providing the iSpinach aptamer and Dr. Simon J. Moore for his help in the cell-free system design. Authors also acknowledge the following funding sources: CONICYT/BC-PhD 72170403 (FM), Biotechnology and Biological Sciences Research Council (BBSRC) [grant number BB/J01446X/1] (MV).

## Supporting Information

**Table S1:**
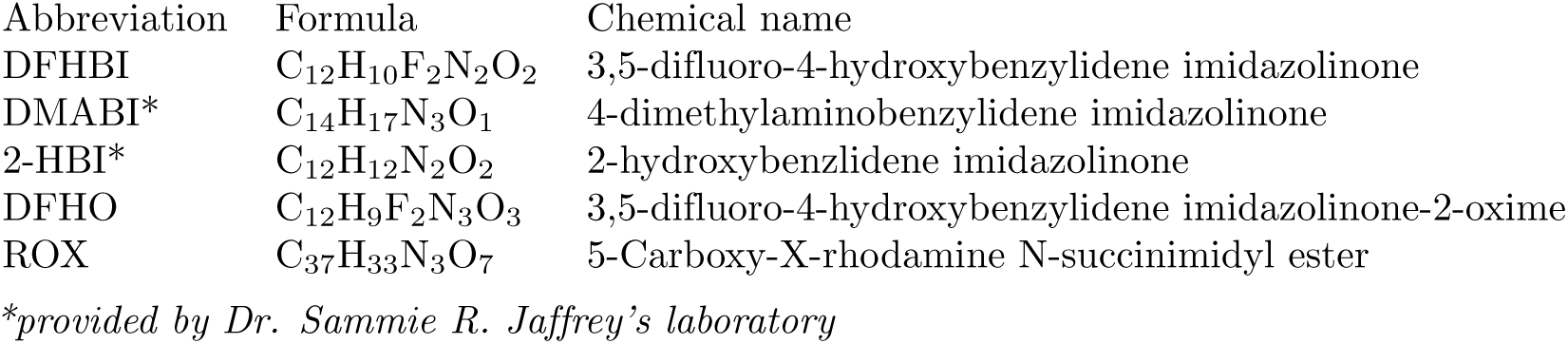
Fluorophores information

| Abbreviation | Formula | Chemical name |
| --- | --- | --- |
| DFHBI | C<sub>12</sub>H<sub>10</sub>F<sub>2</sub>N<sub>2</sub>O<sub>2</sub> | 3,5-difluoro-4-hydroxybenzylidene imidazolinone |
| DMABI* | C<sub>14</sub>H<sub>17</sub>N<sub>3</sub>O<sub>1</sub> | 4-dimethylaminobenzylidene imidazolinone |
| 2-HBI* | C<sub>12</sub>H<sub>12</sub>N<sub>2</sub>O<sub>2</sub> | 2-hydroxybenzylidene imidazolinone |
| DFHO | C<sub>12</sub>H<sub>9</sub>F<sub>2</sub>N<sub>3</sub>O<sub>3</sub> | 3,5-difluoro-4-hydroxybenzylidene imidazolinone-2-oxime |
| ROX | C<sub>37</sub>H<sub>33</sub>N<sub>3</sub>O<sub>7</sub> | 5-Carboxy-X-rhodamine N-succinimidyl ester |

*\*provided by Dr. Sammie R. Jaffrey's laboratory*

**Figure S1:**
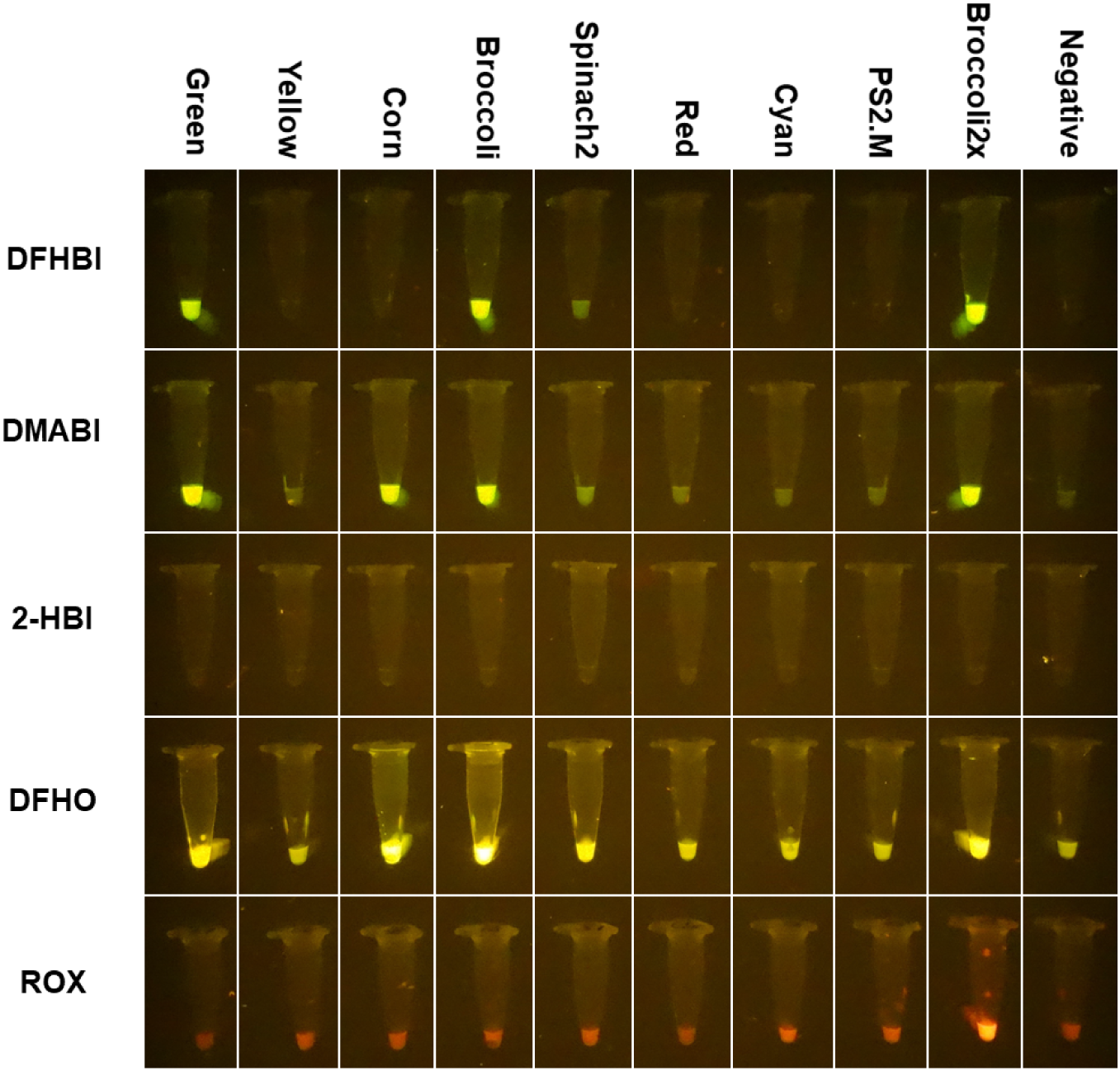
Fluorescent RNA aptamers observed under a blue light/red filter system. Aptamers showing different colours under a blue light/red filter system after measurement 16.5 h and being observed by naked eye without the need for further equipment. Picture taken with a Sony, DSC-RX100 camera and aperture

**Table S2:**
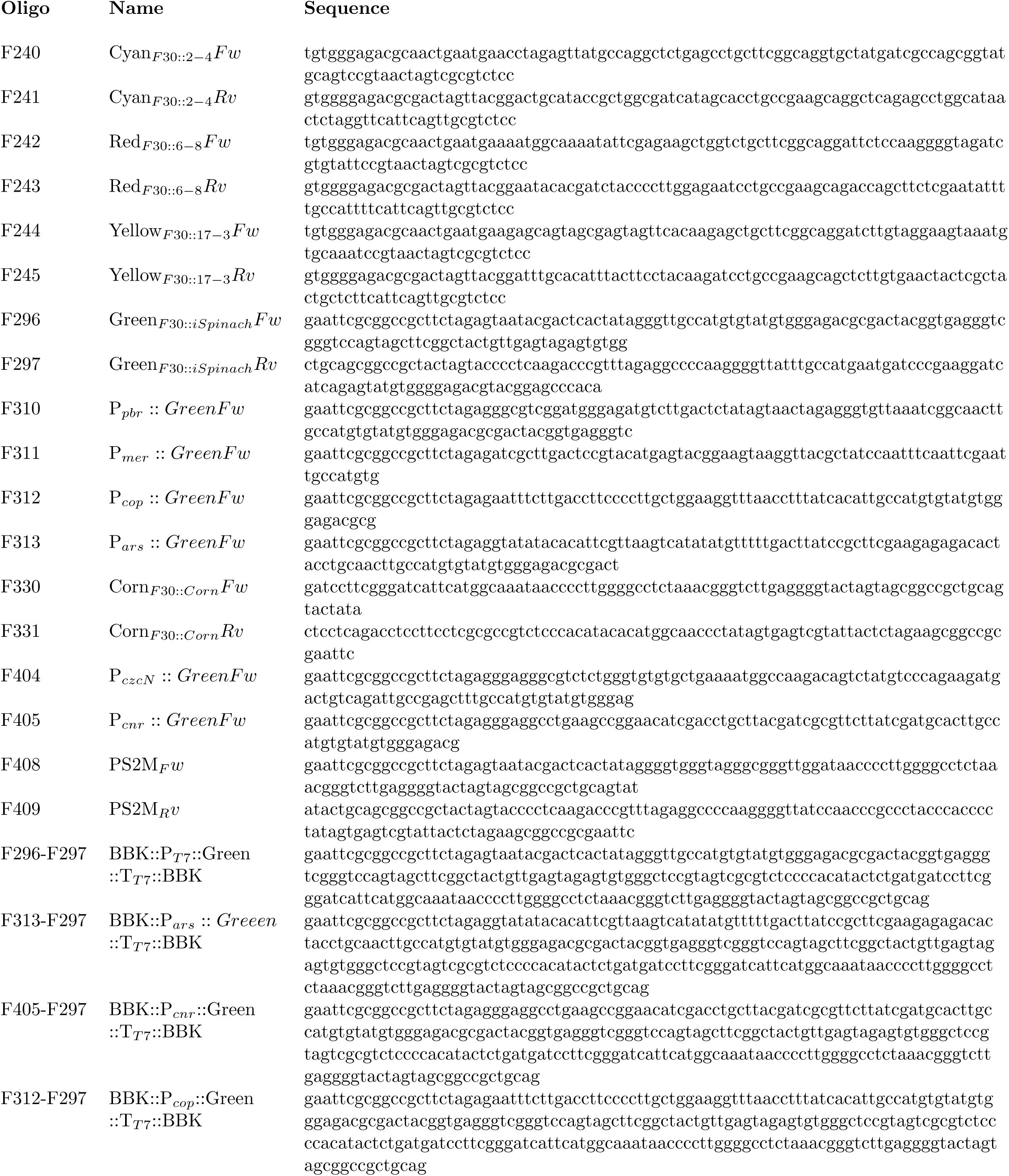
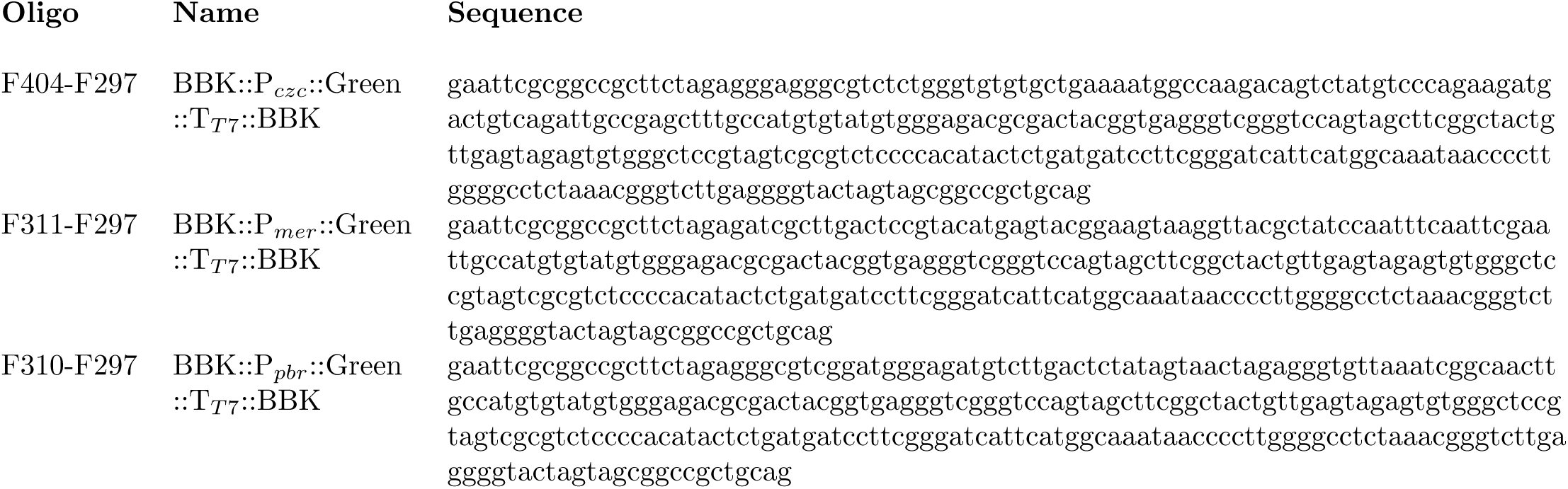
Oligonucleotides used

**Figure S2:**
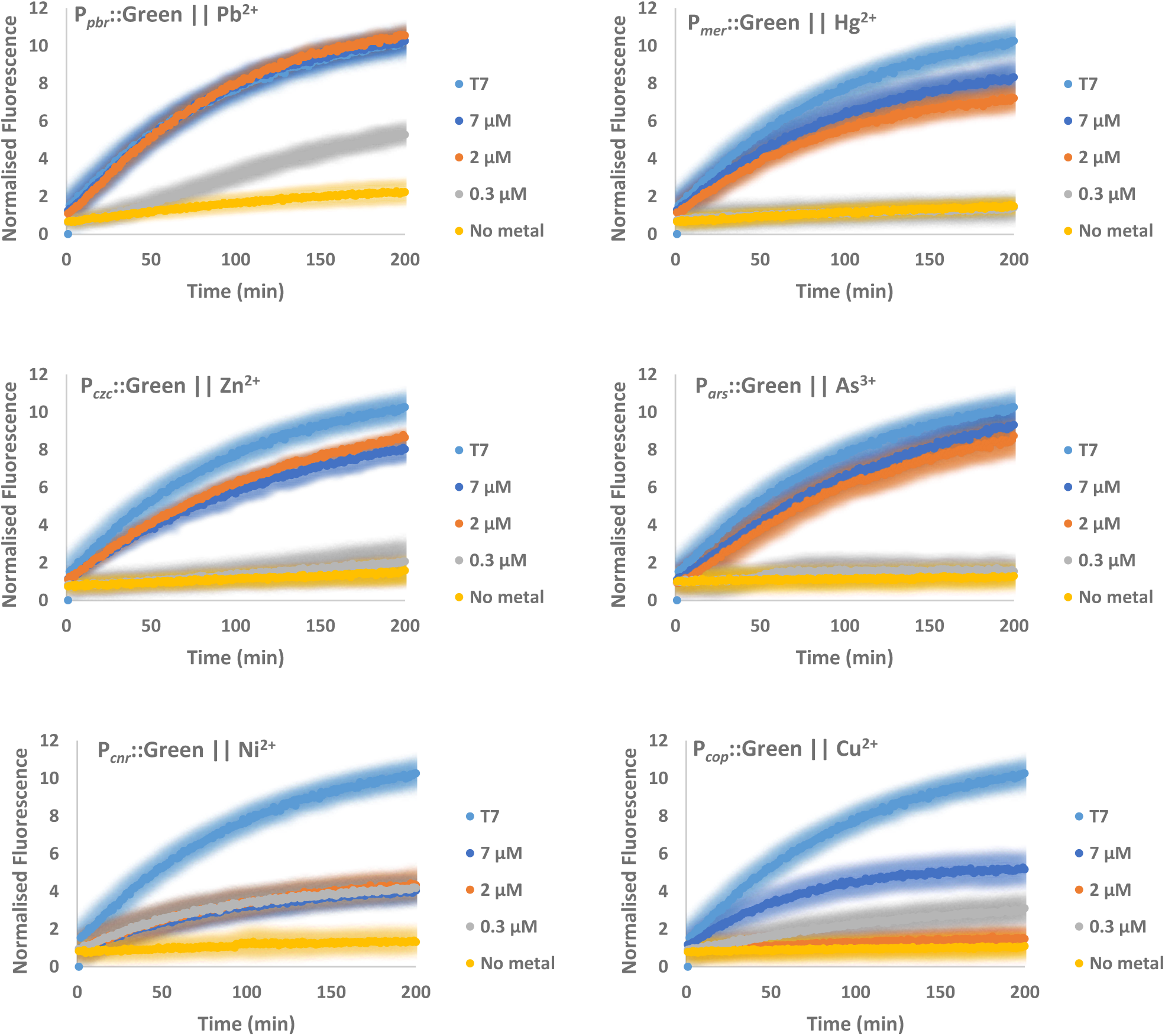
TXO sensors over time with different metals. Coloured shades represent variability of three independent tests.

